# Expelling of *P. falciparum* sporozoites by *Anopheles stephensi* mosquitoes during repeated feeding

**DOI:** 10.1101/2025.11.17.688877

**Authors:** Chiara Andolina, Geert-Jan van Gemert, Jordache Ramjith, Felix Evers, Wouter Graumans, Rianne Stoter, Karina Teelen, Felix JH Hol, Kjerstin Lanke, Teun Bousema

**Affiliations:** Department of Medical Microbiology, Radboud University Nijmegen Medical Centre, Nijmegen, The Netherlands; Institute of Tropical Medicine (ITM), University of Tübingen, Tübingen, Germany; Department of Clinical Research, London School of Hygiene and Tropical Medicine, London, UK

**Keywords:** Interrupted feeding, *Plasmodium falciparum*, sporozoite expelling, artificial skin

## Abstract

It is currently unclear whether repeated mosquito feeding attempts achieve pathogen transmission. We measured *Plasmodium falciparum* sporozoite expelling in *Anopheles stephensi* mosquitoes during two interrupted feeding attempts. The number of expelled sporozoites was positively correlated with salivary gland sporozoite load during the first feeding effort (ρ= 0.36, p= 0.0181) but less so during subsequent feeding (ρ= 0.06, p= 0.6811). The median number of expelled sporozoites was of similar magnitude during the first and second feeding event, with a strong correlation between them (ρ= 0.56, p= 0.0002). We conclude that repeated feeding results in repeated sporozoite inoculation at similar intensity.

## INTRODUCTION

*Plasmodium* parasites are transmitted through the bite of infected female Anopheles mosquitoes. *Anopheles* mosquitoes require a bloodmeal to start oogenesis shortly after adult emergence. Gonotrophic cycles - the time between taking a bloodmeal and laying eggs - take approximately 3 days [1]. Mosquitoes regularly experience interruptions while feeding, due to host defensive movements or failure to access a blood vessel, that can prevent them from completing a full bloodmeal in a single attempt. These interruptions may lead to repeated biting, either on the same host or on multiple hosts during the same gonotrophic cycle. In studies where mosquito bloodmeals were genetically linked to the human blood-source, up to 22% of bloodmeals contained blood from multiple human origins [2, 3]. These meals on multiple human host presumably happened during the same night since DNA is degraded rapidly during bloodmeal digestion [4]. It is unclear whether repeated biting over a short time-window leads to repeated expelling of pathogens. To allow inoculation, sporozoites must move from the secretory cavity to the salivary duct and the dynamics at which this process takes place is poorly understood. Within the salivary duct, sporozoites are organized in parallel or single-file bundles, with only a limited number present there at any given time [5]. Previous studies have shown that inoculum size is highly heterogeneous with a considerable fraction of mosquitos expelling no sporozoites during a given bite despite infected salivary glands [6, 7]. Local sporozoite supply in salivary duct may be limited and it is conceivable that repeated bites over a short time-window may result in few or no sporozoites being inoculated. At the same time, there are indications of repeated transmission events with the same parasite clones at household level, which may result from mosquitoes inoculating sporozoites into multiple hosts during the same night. A study in the Gambia showed that approximately 50% of pairs of children who lived in the same household and were infected with *P. falciparum* carried the same three-locus genotype, whereas only 1% of randomly selected pairs of children were infected with the same parasite genotype [8]. Repeated expelling events by the same mosquito during the same night may explain this observation. Despite the clear implications for malaria transmission dynamics, only one study to date examined how interrupted feeding affects sporozoite ejection. In that study by Ponnudurai and colleagues, *An. stephensi* mosquitoes were fed individually through a mouse skin membrane stretched over human blood. With only 6 mosquitoes used and manual estimation of sporozoite numbers, the study found no evidence for a difference in sporozoite ejection into mouse skin during the first or second feedings [9]. Here, we used a recently developed artificial skin model [6] and sensitive molecular diagnostics to further explore sporozoite expelling by individual *P. falciparum*-infected mosquitoes that were interrupted during feeding.

## METHODS

### *In vitro* cultures of *P. falciparum* gametocytes and mosquito infections

*Plasmodium falciparum* gametocytes (NF54 strain; West-Africa) were cultured in an automated culture system as previously described [10]. Infection feeding procedures are described elsewhere and involved *Anopheles stephensi* mosquitoes, Nijmegen Sind-Kasur strain [6]. Infection burden was assessed 6 days after feeding by counting the number of oocysts on 1% mercurochrome-stained mosquito midguts. A second uninfected bloodmeal was provided 7 days post infection (PI) to synchronize oocysts development [11].

### Mosquito feeding experiments

Fifteen days post infection, individual mosquitoes were collected in small acrylic cages (5x5x7cm), covered with netting material on the top and bottom. Mosquitoes were starved 4 hours prior to feeding experiments. Artificial skin (INTEGRA® dermal substitute) was secured to a glass membrane feeder connected to a heated circulating water bath set to 39°C; 100µl of human blood was pipetted onto the artificial skin’s surface (∼ 1.4 cm^2^), and spread evenly for mosquito feeding [6]. The experiments were performed in a secure insectary with the room set at 21°C and 80% relative humidity, to ensure a good heat stimulus for feeding. For the first feeding, the feeder was placed on top of the cage, allowing the mosquito to probe through the membrane for a maximum of one minute or until a partial bloodmeal was observed. Mosquitoes were closely inspected during probing and feeding. Feeding was interrupted once a partial bloodmeal was observed in the abdomen or if the mosquito was in contact with the membrane for approximately 60 seconds. After the first artificial skin was removed, the same mosquito was allowed to probe on a second artificial skin until it had taken a full bloodmeal. After the two feedings, mosquitoes were dissected under a stereo microscope and salivary glands were collected in 1,5mL Eppendorf tubes containing 180µl oocyst lysis buffer (NaCl 0.1M: EDTA 25mM: TRIS-HCl 10mM). A scalpel (Dalhausen präzisa plus, no 11) was used to cut the artificial skin above the rubber band around the entire feeder. The artificial skin was transferred with a DNA-free tweezer to a 1,5mL Eppendorf tube containing 180µl oocyst buffer (NaCl 0.1M: EDTA 25mM: TRIS-HCl. 493. 10mM), and stored at -70°C. *P. falciparum* sporozoites were quantified by COX1-qPCR as described elsewhere [6].

### Statistical analysis

Statistical analyses were performed in R (version 4.3.2). Analyses excluded mosquitoes without detectable salivary gland sporozoites. Expelling prevalence was modelled using a logistic regression model with total sporozoite load category and feeding round as fixed factors. Model estimates with 95% confidence intervals were plotted to visualise expelling probabilities (Figure 1A). Correlations between (i) salivary gland sporozoite load and expelled sporozoites in Feed 1 and Feed 2, (ii) expelled sporozoites between Feed 1 and Feed 2, and (iii) the corresponding fractions expelled were assessed using Spearman’s rank correlation (ρ). Paired differences in expelled numbers between Feed 1 and Feed 2 were evaluated using Wilcoxon signed-rank tests. A two-sample test for equality of proportions compared the fraction of mosquitoes expelling sporozoites in Feed 1 and Feed 2. All tests were two-sided with α = 0.05.

**Figure 1.**
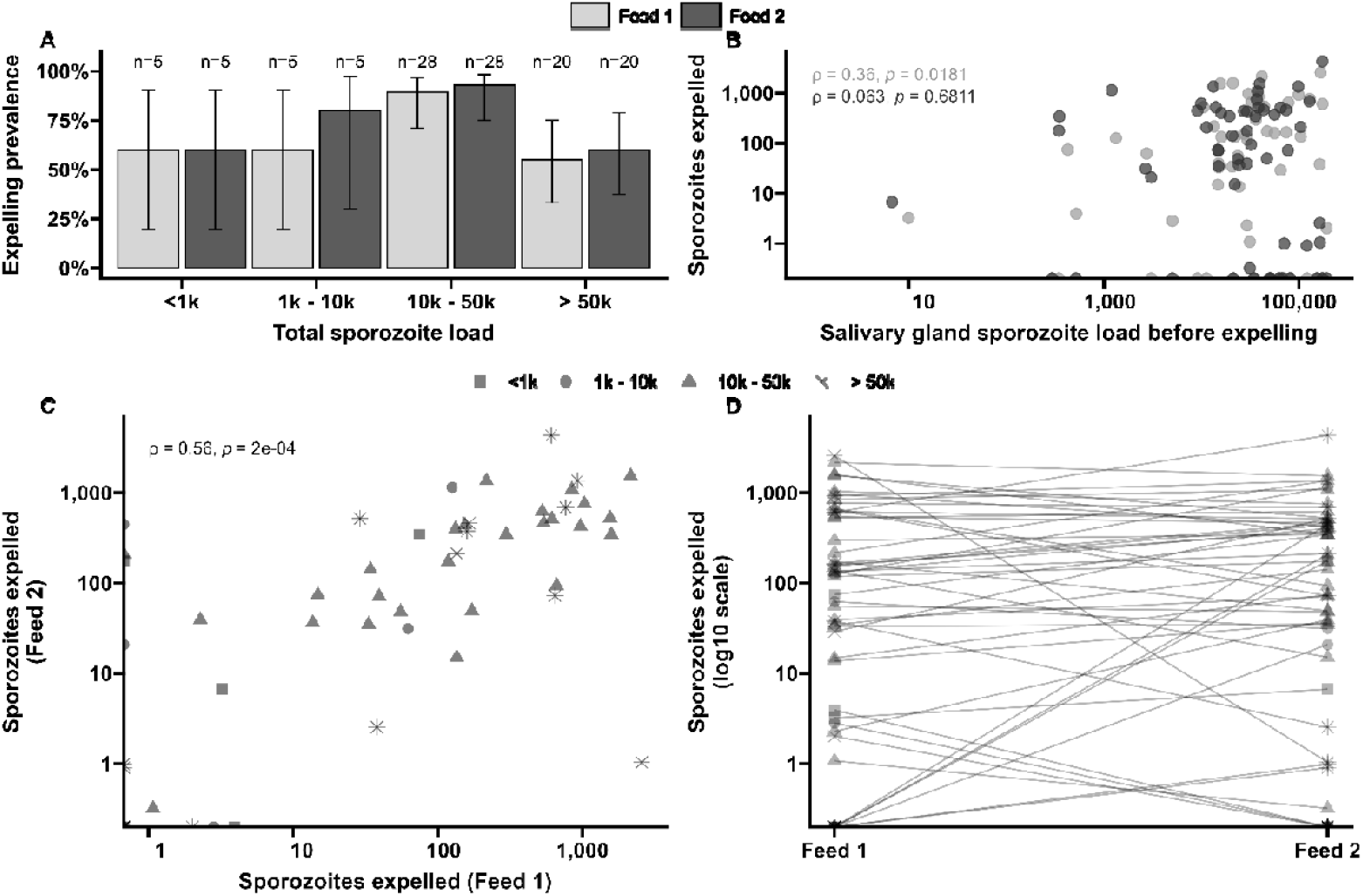
(**A**) Total sporozoite load in artificial skin and salivary glands (x-axis) were binned by infection load <1k, 1k-10k, 10-50k, >50k and plotted against the proportion of mosquitoes (%) that expelled sporozoites (y-axis) as estimated from an additive logistic regression model with factors Feed 1 and Feed 2 categories. (**B**) Salivary gland sporozoite load before expelling (sporozoites in salivary glands + sporozoites expelled, x-axis) in relation to the number of expelled sporozoites (y-axis) for the Feed 1: ρ = 0.36 (p=0.0181) and for Feed 2: ρ = 0.063 (p=0.681). Sporozoite numbers are shown on a log 10 scale. (**C**) The correlation between the number of sporozoites expelled in Feed 1 and Feed 2 among mosquitoes that expelled sporozoites in both feeds. Each symbol represents an infected mosquito that fed twice. Different symbols represent mosquitoes from distinct infection load groups. (**D**) The total number of sporozoites expelled during both the first and second feeds across all categories. Each pair of points that is connected with a line represents an individual mosquito. Mosquitoes that never expelled sporozoites are not included in this graph.

### Ethics declarations

Experiments with *in vitro* cultured parasites and *An. stephensi* mosquitoes at Radboud university medical center were conducted following approval from the Radboud University Experimental Animal Ethical Committee (RUDEC 2009-019, RUDEC 2009-225).

## RESULTS

Four independent batches of 20 mosquitoes with 100% oocyst prevalence, assessed on day 6 post-infection, were used for experiments (median number of oocysts=14; IQR=10-20). Individual mosquitoes were collected in small cages and allowed to probe on artificial skin. After approximately one minute since the mosquito came in contact with the artificial skin and a partial bloodmeal was observed, feeding was disturbed and a second artificial skin was offered to allow the mosquito to feed until repletion. A total of 60 mosquitoes was used for the experiments. Of these, 98% (58/60) had sporozoites detected in either the salivary gland or the skin. One mosquito had no detectable salivary gland sporozoites, yet expelled 3 and 7 sporozoites during the first and second feedings, respectively. This mosquito was excluded based on the negative salivary gland and very low skin sporozoite estimates. Finally, one mosquito neither had salivary gland sporozoites or expelled sporozoites and was concluded to be non-infected and thus excluded. In the first feed, 72% (42/58) of mosquitoes expelled sporozoites, while in the second feed, 77% (45/58) of mosquitoes expelled sporozoites (p=0.6680) Among those mosquitoes that did not expel in the first feed (n=16), 63% (10/16) also did not expel in the second feed. Conversely, among those that expelled in the first feed (n=42), only 7% (3/42) did not expel in the second feed. Generally, 67% (39/58) of mosquitoes expelled sporozoites in both feeds, 83% (48/58) expelled in at least one of the feeds, and 17% (10/58) expelled in neither. Among mosquitoes that expelled sporozoites in both feeds (N=39), the median number of expelled sporozoites in the first feed was 159 (IQR= 47.3 – 656.2), while in the second feed it was 346 (IQR= 48.8 – 521.4) (p=0.7500). Mosquitoes that failed to expel in both feeds (N=10) had a median total sporozoite salivary gland load of 56,962 (38, 503 – 77,707) (**Table 1**) (**Figure 1C**), which is non-significantly (p=0.2990) higher than the load in mosquitoes that expelled in both feeds (n=39; median load 30284; IQR 15,834-51,745). To investigate the relationship between sporozoite expelling and infection burden, mosquitoes were binned into four categories based on total sporozoite load (the total number of residual salivary gland sporozoites plus the total number of expelled sporozoites) (**Figure 1A**) [6]. We observed a weak but statistically significant association between salivary gland sporozoite load before expelling and sporozoites expelled in the first feed (Spearman’s correlation coefficient ρ= 0.36, p= 0.0181; N=42) (**Figure 1B**), indicating that mosquitoes with a higher salivary gland sporozoites load tend to expel more sporozoites during the first feed. In the second feed, no statistically significant association was observed (ρ= 0.06, p= 0.6811; N=45).

**Table 1.**
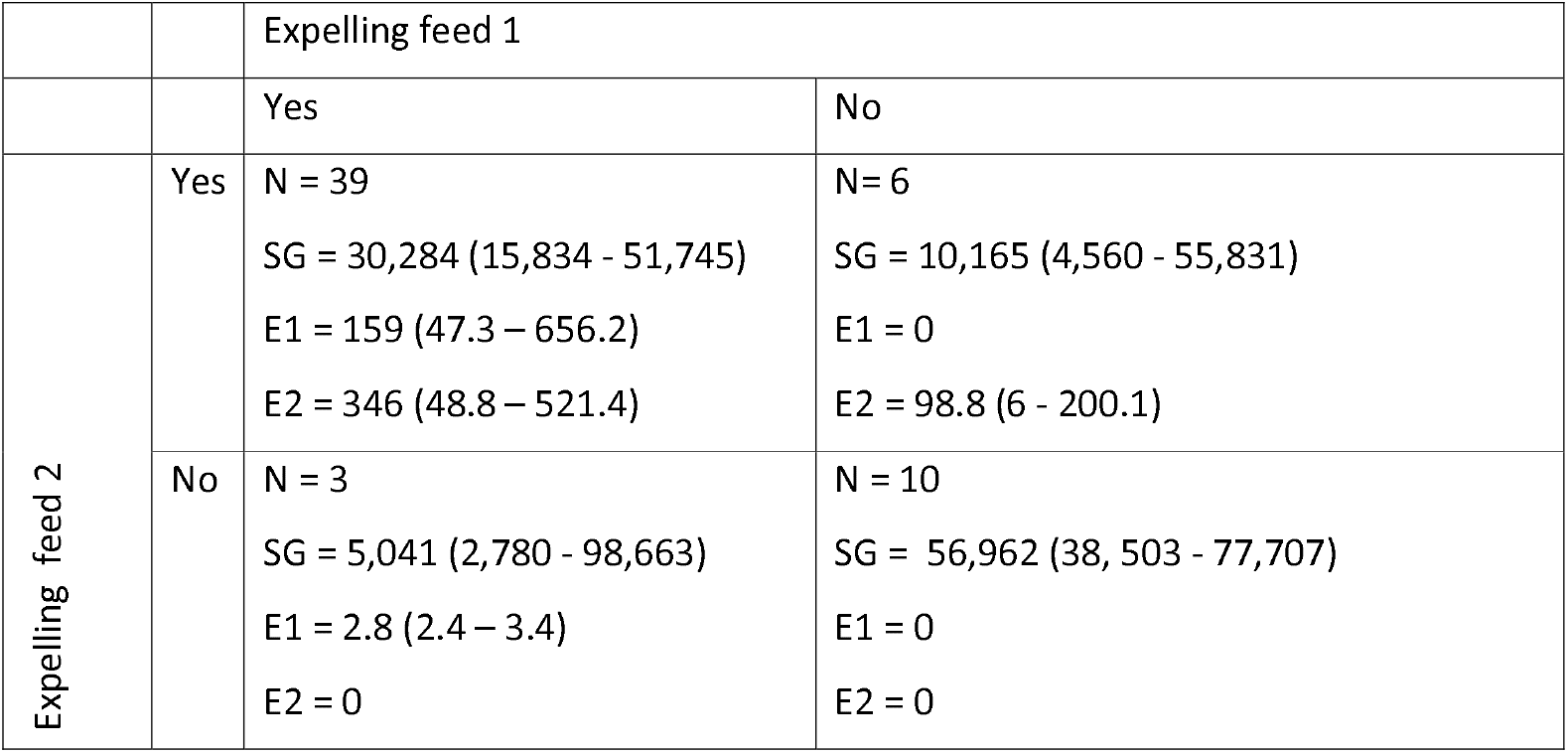
Median Sporozoites loads (IQR) in mosquitoes after interrupted feedings. Cells display the number of mosquitoes that expelled in feed 1 and/or feed 2 and characteristics of sporozoite load and inoculum. Each cell contains: the number of mosquitoes in this category (N), the salivary gland load (SG) as median with interquartile range, the number of expelled sporozoites in the first feeding (E1) and second feeding (E2).

|  |  | Expelling feed 1 |  |
| --- | --- | --- | --- |
|  |  | Yes | No |
| Expelling feed 2 | Yes | N = 39<br>SG = 30,284 (15,834 - 51,745)<br>E1 = 159 (47.3 – 656.2)<br>E2 = 346 (48.8 – 521.4) | N= 6<br>SG = 10,165 (4,560 - 55,831)<br>E1 = 0<br>E2 = 98.8 (6 - 200.1) |
|  | No | N = 3<br>SG = 5,041 (2,780 - 98,663)<br>E1 = 2.8 (2.4 – 3.4)<br>E2 = 0 | N = 10<br>SG = 56,962 (38, 503 - 77,707)<br>E1 = 0<br>E2 = 0 |

We observed a strong association between sporozoites expelled in the first feed and in the second feed ρ= 0.56, p= 0.0002 (**Figure 1C**). This indicates a level of consistency in the number of sporozoites expelled that was also observed when sporozoite expelling was quantified as the fraction of the total sporozoite load (ρ= 0.73, p= 0.0001 **Supplementary Figure 3 A-B**).

## DISCUSSION

The present study aimed at quantifying the number of sporozoites expelled during interrupted feedings by employing sensitive molecular techniques combined with individual mosquito probing and feeding experiments on artificial skin. We observed that 80% of infected mosquitoes expelled sporozoites in at least one of two feeding attempts, with 65% expelling in both. Moreover, we observe a significant correlation between the number of sporozoites expelled in the first and second feeding attempt and no evidence for declining inoculum size upon repeated feeding.

Before each gonotrophic cycle a mosquito may take one or more bloodmeals, with the frequency of feeding being associated with *Anopheles* species [12, 13] and potentially whether a mosquito is infected with *Plasmodium* [12]. Additionally, low levels of salivary apyrase, an anti-platelet aggregation enzyme, may lead to prolonged probing time and incomplete bloodmeals, plausibly increasing the biting rate [14]. The consequences of repeated feeding on the likelihood that malaria parasites are transmitted remain largely unstudied. An early study by Rosenberg, in which mosquitoes were attached to a glass slide and forced to salivate into mineral oil, indicated that the majority of *P. falciparum* sporozoites are expelled at the onset of salivation and longer salivation has diminishing returns on transmission potential [15]. In contrast, for *P. berghei* it was reported that transmission potential may persist throughout multiple probing attempts, rather than being limited to the initial feed [16].

Given that time is required for sporozoites to enter the duct, we hypothesized that repeated feeding over a short time window would lead to a smaller chance of expelling. Our study was specifically designed to test this hypothesis and quantify sporozoites expelled in subsequent feeding attempts. For this, we used a skin mimic that allows mosquitoes to take a bloodmeal and was previously used with both *An. stephensi* and *An. coluzzii* mosquitoes [6]. When comparing the absolute number of sporozoites, or the fraction of the total sporozoite load, that is expelled before and after interrupted feeding, we observed no evidence of a lower sporozoite inoculum in the second feeding attempt. Instead, we observed a strong correlation between the number of sporozoites expelled during the first and second meal. Mosquitoes capable of expelling sporozoites during one feeding attempt were more likely to also expel sporozoites in a subsequent attempt. This is in line with a limited number of experiments where single infected mosquitoes were interrupted during feedings through a mouse skin membrane, and sporozoites were counted using fluorescence microscopy, which showed no evidence of depletion [9]. Our data suggest that even mosquitoes with a relatively low infection burden (10k-50k sporozoites in the salivary glands, reflecting ∼2 - 10 oocysts [6]), can effectively transmit malaria to multiple hosts.

Intriguingly, repeated biting may be increased for infected mosquitoes. A study in Tanzania investigating whether *An. gambiae* infected with *P. falciparum* sporozoites fed on more hosts than uninfected mosquitoes, found that 22% of infected mosquitoes took multiple bloodmeals, compared to only 10% of uninfected mosquitoes [12]. Interestingly, we observed that mosquitoes with high sporozoites load (median 56,962) failed to expel sporozoites in both feedings. Similar observations have been made in mosquitoes with very high number of sporozoites in the salivary glands, where only a small number of sporozoites were expelled [6, 7, 17]. The complex architecture of salivary glands and potential damage to this architecture following high intensity infection, might influence expelling efficiency [17].

Our study has several limitations. First, we had a relatively modest sample size that reflected the laborious nature of experiments with individual mosquitoes. Whilst this sample size was insufficient to detect subtle differences in sporozoite inoculum size between feeding attempts, the fact that we observed a numerically higher inoculum size in the second feeding attempt gives us confidence that we can reject our hypothesis that the number of expelled sporozoites declines during repeated feeding. A second limitation is that the duration of the second feeding attempt was not standardized or forced to be identical to the first feeding attempt. The second feeding may thus be longer and it is possible, albeit not proven, that this would associate with a higher inoculum size [5, 7]. Heterogeneity in mosquito behaviour and difficulties in disentangling the subtleties of mosquito behaviour in our assay (e.g. directly observing the stylet and confirming effective probing or quantifying the duration of engorgement) makes it unrealistic to achieve higher precision with our set-up. Lastly, our sporozoite load was higher than typically observed in natural infections. Future studies might examine whether our findings can be extrapolated to mosquitoes with low infection burdens.

We conclude that repeated feeding attempts frequently lead to repeated sporozoite inoculation and provide further evidence that mosquito salivary gland load is a relevant determinant of inoculum size.

## Supplementary data

Supplementary materials are available at The Journal of Infectious Diseases online (http://jid.oxfordjournals.org/). Supplementary materials consist of data provided by the author that are published to benefit the reader. The posted materials are not copyedited. The contents of all supplementary data are the sole responsibility of the authors. Questions or messages regarding errors should be addressed to the author.

## Acknowledgments

We would like to thank all the entomology staff at Radboudumc, Laura Pelser-Posthumus, Astrid Pouwelsen, Jolanda Klaassen, and Jacqueline Kuhnen for all mosquito husbandry; Claudia Bin for all the tester units of Integra Dermal Regeneration Template.

## Supplementary

**Supplementary Table 1:**
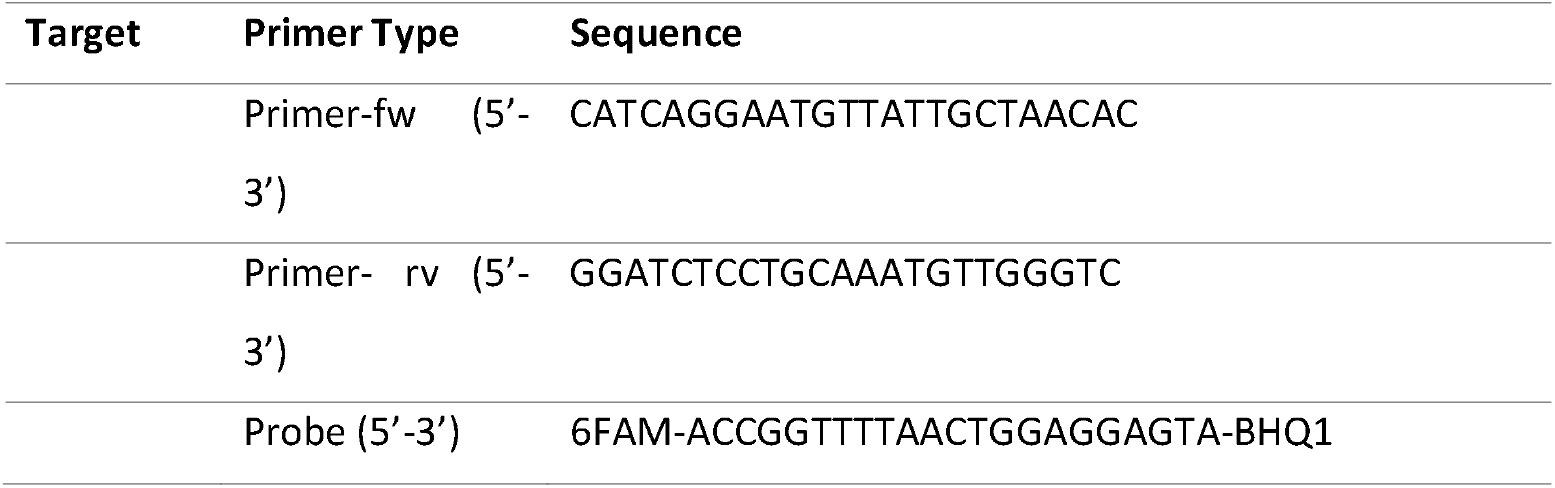
Primer and probe sequence used in COX-I sporozoite qPCR.

**Supplementary Figure 1.**
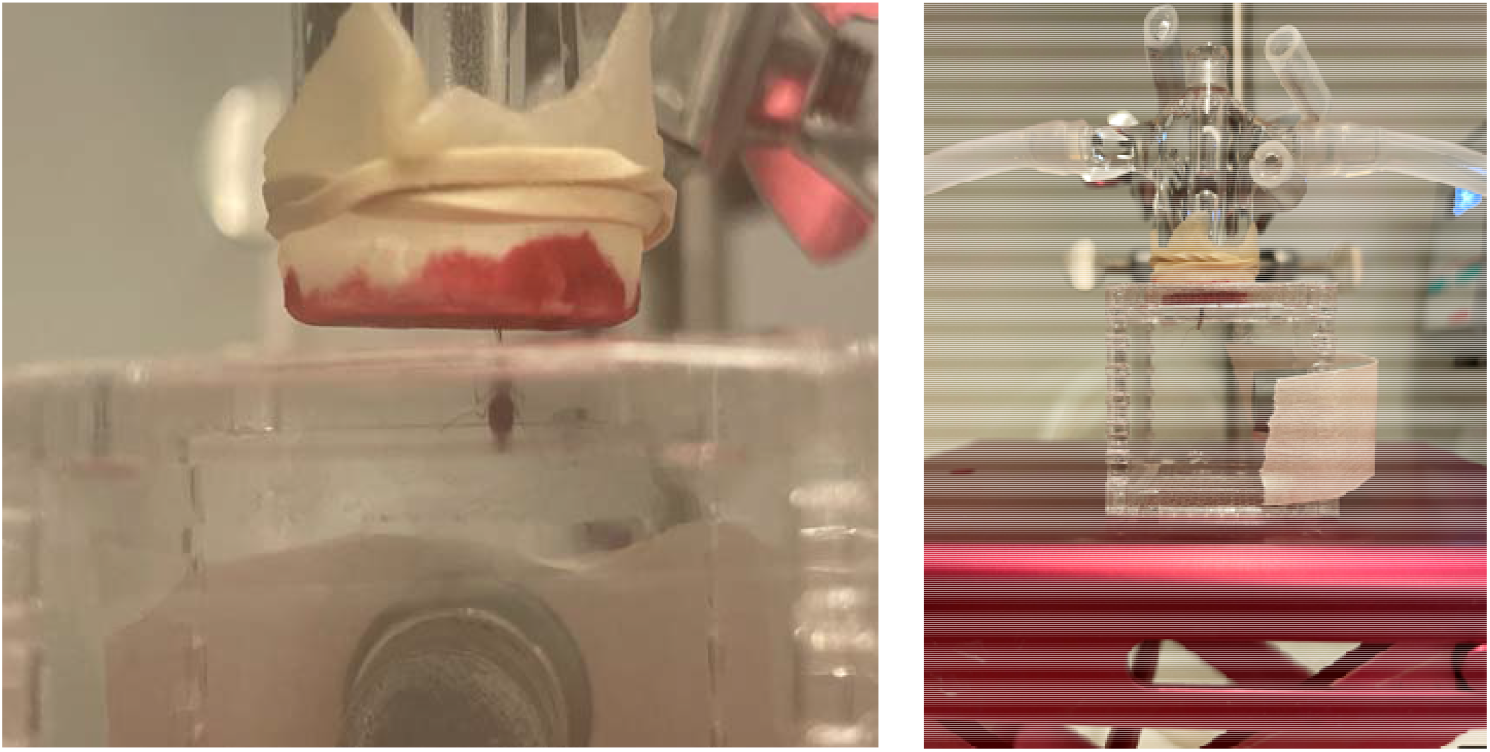
*An. stephensi* mosquito feeding on artificial skin covering a glass feeder connected to a circulating water bath set at 37C.

**Supplementary Figure 2.**
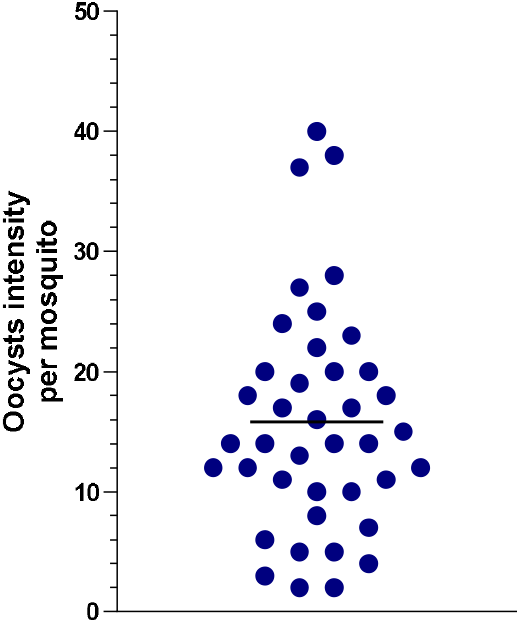
Scatter plot of oocysts distribution on day 6 PI for 4 experimental infections. The bar shows the mean.

**Supplementary Figure 3.**
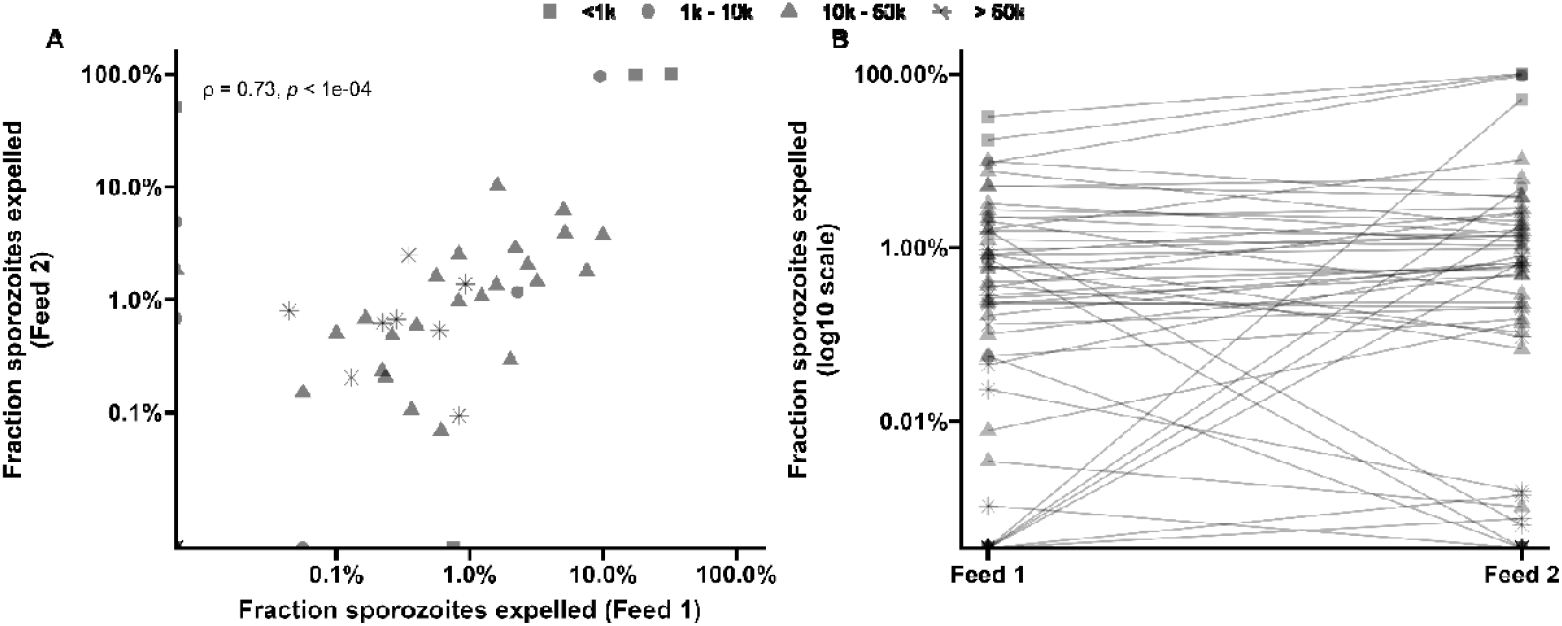
Correlation between the fraction of sporozoites expelled during the first and second blood feeds. **A**. Each point represents an individual mosquito, with the fraction of sporozoites expelled during Feed 1 (x-axis) plotted against the fraction expelled during Feed 2 (y-axis), both shown on a logarithmic scale. Different symbols (triangles, squares, asterisks, circles) correspond to distinct infection groups. **B**. Each connected line represents one mosquito, showing the fraction of sporozoites expelled during Feed 1 and Feed 2 on a log10 scale. Symbols denote different infection groups, as in Panel A.

